# Heat-induced secondary dormancy contributes to local adaptation in Arabidopsis thaliana

**DOI:** 10.1101/2025.02.20.638146

**Authors:** Nhu Loc Thuy Tran, Tahir Ali, Gregor Schmitz, Juliette de Meaux

## Abstract

Seeds should not germinate in conditions unsuitable for seedling growth. Dormancy, which allows seeds to remain inactive in an environment that would otherwise enable germination, helps optimize the timing of germination. Primary dormancy, developed during seed maturation on the parent plant, prevents immediate germination post-dispersal, regardless of external conditions. Secondary dormancy, however, is triggered post-dispersal when seeds face unfavorable conditions, enabling them to re-enter dormancy even if initially non-dormant. This mechanism allows seeds to fine-tune germination according to environmental conditions. In this study, we examined the role of heat-induced secondary dormancy in local adaptation by analyzing natural variations within 361 *Arabidopsis thaliana* accessions from across Europe. We discovered that secondary dormancy acquisition varies with primary dormancy levels and after-ripening. Both primary and heat-induced secondary dormancy exhibited adaptive clines along temperature and precipitation gradients, with secondary dormancy showing a steeper cline, indicating its significant role in local adaptation. Using species distribution models, we predicted that genotypes with high secondary dormancy would show greater resilience to future climate changes. Additionally, we identified specific genomic regions controlling secondary dormancy levels, including a novel candidate gene for secondary dormancy variation. Our findings show that secondary dormancy is a complex adaptive mechanism and a predominant contributor to the dormancy trait syndrome that favors plant survival in habitats exposed to harsh summers.

**Summary statement:** Secondary dormancy induced by heat exposure allows seed to adjust their germination strategies to the environment. This study shows that heat-induced secondary dormancy in Arabidopsis thaliana depends on levels of primary dormancy. Its covariance with climatic parameter indicates that it can contribute to population resilience to climate change. This study further identifies the specific genetics underlying the ability to induce dormancy after dispersal.

## Introduction

Coping with environmental changes is key for living organisms to survive. In nature, plants perceive and adjust to their environment by synchronizing their life stages to the seasons (Blackman, 2017; Heggie & Halliday, 2005; Lamers et al., 2020). However, unpredictable changes in seasonal patterns, such as shifts in temperature or rainfall, can disrupt the timing of biological events and the interactions between organisms and their environment, a process often referred to as ecological dynamics. (Hamann et al., 2021; Sohindji et al., 2020). One critical aspect of these interactions is the seasonal timing of germination, which is the first key life-history transition in a plant’s life cycle (Baskin and Baskin, 1998). The timing of germination not only determines the conditions under which seedling establishment and growth occur, but also influences post-germination traits, such as vegetative growth and flowering time (Burghardt et al., 2015; Donohue et al, 2010; Wagmann et al., 2012; Zacchello et al., 2020).

Germination timing is regulated by seed dormancy, a mechanism that prevents seed germination under unfavorable conditions (Chahtane et al., 2017; Lamont & Pausas 2023). Like all phenological traits shaping plant life histories, dormancy has evolved in plants to align with their habitats. Global analyses show that seed dormancy levels generally vary along major climatic gradients. Where summer precipitation is regular and temperatures are predictable, such as in subtropical or northern habitats, plant species rarely produce dormant seeds (Zhang et al. 2022). Conversely, species growing in habitats experiencing unpredictable summers (unusually warm or dry) tend to have more dormant seeds (Zhang et al. 2022). For example, the survival of early spring germinants could be compromised if germination is followed by a hot and dry summer. In such conditions, delaying germination until after the predicted challenging season increases seedling survival and fitness (Chiang et al., 2013; Donohue et al., 2010).

Dormancy can vary between populations of a species, as well as among different species (Willis et al., 2014). Two main types of dormancy exist: primary and secondary. Primary dormancy is established while the seed develops on the mother plant; secondary dormancy is established after seed dispersal, upon seed exposure to unfavorable conditions (Baskin and Baskin, 1998; Soltani et al., 2019). This capacity to respond dynamically to environmental cues provides additional flexibility beyond primary dormancy and can be particularly advantageous in unpredictable climates. Variation in primary dormancy has been documented extensively in the model plant Arabidopsis thaliana (Bentsink et al. 2006; Debieu et al. 2013; Footitt et al., 2011, 2015; Martel et al., 2018), but also within other species growing in a wide range of climatic zones, such as in Korean milkweeds, Asclepias (Kaye et al., 2018) or among species of the Fabaceae family (Wyse and Dickie, 2017). In *A. thaliana*, genetic variation in primary dormancy contributes to local adaptation (Kronholm et al. 2012; Chiang et al. 2013, Kerdaffrec & Nordborg, 2017). The cline of genetic variation for seed dormancy observed in *A. thaliana* follows the pattern observed across plant species: higher levels of primary dormancy are observed in regions where the growing season is long, but summer can be very dry (Postma and Ågren, 2022; Klupczyńska and Pawłowski, 2022).

In nature, temperature fluctuations shape plant perception of seasonal change (Bujs, 2020; Chahtane et al., 2017; Klupczyńska and Pawłowski, 2021). Maternal environments, particularly, summer temperature and summer precipitation, can have strong impacts on the magnitude of seed dormancy (Kerdaffrec & Nordborg, 2017; Zacchello et al., 2020; Chiang et al., 2013; Coughlan et al., 2017). Studies in *A. thaliana* and other common model species, such as oilseed rape and oats, have shown that a so-called secondary dormancy can be induced after seed dispersal by a few days of exposure to high or low temperatures in experimental and field settings (Buijs et al., 2020; Footitt et al., 2011, 2015; Martel et al., 2018; Malavert et al., 2017; Pawłowski et al., 2020; Gulden et al., 2004, Ņečajeva, et al, 2021). Late summer temperatures have been confirmed to strongly delay germination of *A. thaliana* seeds in nature (Schmitz et al. 2023).

Despite the obvious ecological implications of secondary dormancy, its genetic variation and adaptive relevance is poorly understood. Ecologically, cold-induced dormancy has been documented in species like *Butia odorata* (Schlindwein et al., 2019), Rosaceae, and poplars (Yamane et al., 2021), as well as various species such as Turkish pine *Pinus brutia* Ten., black adler *Alnus glutinosa* (L.) among others (Klupczyńska and Pawłowski, 2021). In *A. thaliana*, primary dormancy and cold-induced secondary dormancy correlate (Debieu et al. 2013, Martinez-Berdeja et al. 2020). The correlation was positive in northern latitudes, but changed sign in southern latitudes (Debieu et al., 2013). Cold-induced dormancy could thus form part of some of the adaptive trait syndrome that evolved to diversify ecological strategies along the species range (Takou et al., 2018, Martinez-Berdeja et al. 2020, Exposito-Alonso et al. 2020).

Whether dormancy induced by heat after dispersal complements these traits is unknown. Heat-induced secondary dormancy has been documented in the pioneering work of Bouwmeester and Karssen (1993) and Baskin and Baskin (1997): seeds from summer annuals acquire secondary dormancy after exposure to high summer temperatures. Secondary dormancy has been documented, for example, for the tropical species Comanthera bisulcata and Syngonanthus verticillatus as well as the mediterraneaan Cistus species (Duarte and Garcia, 2015; Zomer at al. 2022). It may also be centrally relevant for biodiversity conservation in fire-prone environments (Cuena Lombraña et al., 2024). In *A. thaliana*, exposure to warm temperatures at the end of summer led to delayed germination and altered expression of genetic variation in flowering time and fitness (Schmitz et al. 2023). It is, therefore, crucial to determine whether variation in secondary dormancy sustains adaptation to local environmental conditions.

Additionally, our understanding of the molecular basis of secondary dormancy remains rudimentary (Iwasaki et al., 2022; Bujs, 2020; Gianetti, 2023). Even though primary and secondary dormancy clearly differ according to the conditions under which they are induced, the mechanisms that govern their respective establishment are not elucidated (Iwasaki et al. 2022; Bujis, 2020). The DELAY-OF-GERMINATION-1 gene (DOG1) is known as a major regulator of natural variation in primary seed dormancy (Bentsink et al 2006, Chiang et al. 2013) and is believed to condition the ability of the seed to respond to abscisic acid (ABA) signals that can arrest germination (Iwasaki et al., 2022). The activity of DOG1 decreases in dry seed in a process called after-ripening, which is not yet fully understood but seems to depend on redox processes (Nee et al., 2017). Interestingly, genotypes that did not become dormant after prolonged cold exposure tended to have weak primary dormancy and associated with a non-functional *DOG1* haplotype (Martinez-Berdeja et al. 2020). Levels of heat-induced secondary dormancy also depend partially on residual primary dormancy (Bujis, 2020; Coughlan et al., 2017). Primary and secondary dormancy may therefore be regulated by similar pathways.

In this study, we quantified natural variation in heat-induced seed dormancy and used environmental associations to document its adaptive relevance in *A. thaliana*. Using comparative niche modeling, we report that strong dormancy genotypes are likely to be more resilient to future changes in climate. Our study confirms that secondary dormancy variation changes with primary dormancy; however, genome-wide association studies show that the variation in secondary dormancy is under the control of specific genetic variants. This study underscores the ecological significance and intricate molecular basis of secondary seed dormancy and highlights its crucial role in plant adaptation.

## Materials and Methods

### Seed material

The samples used in this study originated from a collection of 361 accessions across Europe (Suppl. Figure S1). Genotype and accession information was obtained from the 1001 genome database (1001genomes.org; 1001 Genomes Consortium, 2016) as well as from Wieters et al. (2021) (Suppl. Table S1). Seed material was amplified at University of Cologne: plants were grown in growth chambers (Dixell, Germany) under long day condition 16:8 (hours) light: dark at 20^0^C (day): 18^0^C (night) (hereafter referred to as “standard condition”). Plants were vernalized for 4-6 weeks, depending on their genotype, to ensure the required vernalization period was met. As a result, most plants produced seeds around the same time, with a maximum difference of one week. Mature dry seeds were harvested and packed in paper bags; these seeds were stored at room temperature for approximately six months post-harvest and subsequently transferred to 4°C for long-term storage. For each genotype, we harvested one seed batch of around 1000 seeds, from which we performed dormancy experiments 6 months, one year, and two years after harvesting, i.e. May 2022 – Trial 1, December 2022 – Trial 2, and December 2023 – Trial 3. Each experiment used 50-200 seeds. Due to practical limitations, the first Trial 1 used a set of 295 accessions; Trial 2￼￼ and the Trial 3, a set of 344 accessions.

### Germination test

50-200 seeds per genotype were sown on wet filter paper in 12-well plates in a randomized design. There were three incubation treatments, namely, “control,” “primary dormancy,” and “secondary dormancy” (Figure 1). As a viability control, seeds were pre-incubated at 4^0^C for three days and moved to standard condition. To assess primary dormancy, we put seeds for germination without pre-treatment. To assess secondary dormancy, seeds were pre-incubated at 4^0^C in the dark for three days to release primary dormancy, then exposed to 37^°^C for four days, and finally moved to standard condition for germination. Germination rate was assessed after seven days, with radicle protrusion serving as the criterion for germination. The number of germinated and non-germinated seeds was recorded by carefully examining radicle emergence under a stereomicroscope. Approximately 50-200 seeds were placed in each well and carefully counted.

**Figure 1.**
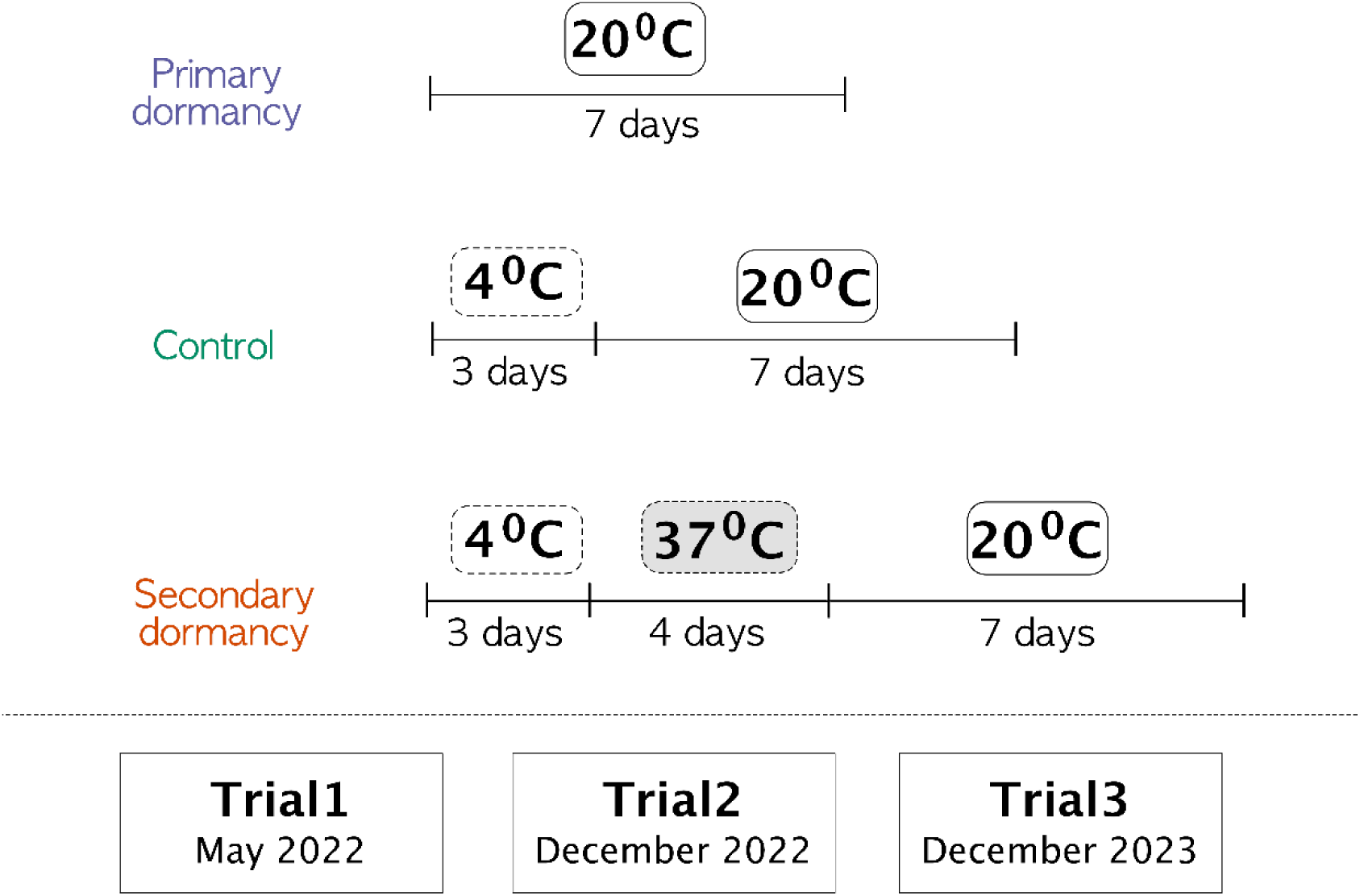
Overview of the experimental design for studying primary and heat-induced secondary dormancy. This schematic illustrates the three treatments used to assess germination behavior and seed dormancy: primary dormancy, secondary dormancy, and control. For the control treatment, seeds were stratified at 4°C for three days to release any dormancy before being exposed to germination conditions. In the primary dormancy treatment, seeds were tested directly without any pre-treatment to assess the baseline dormancy established during seed maturation. For the secondary dormancy treatment, seeds underwent stratification at 4°C for three days to release dormancy, followed by a heat stress of 37°C for four days to induce secondary dormancy. Approximately 50–100 seeds were used per treatment in each genotype. Germinated and non-germinated seeds were counted after a seven-day germination period under long-day conditions (16 hours light, 20°C). Germination rates, calculated based on the proportion of germinated seeds, were used in subsequent analyses.

### Bioclimatic data

Climatic variables were obtained from the WorldClim database (worldclim.org/version2) using the worldclim_global function (R/geoData package, Hijmans et al., 2022). Last Glacial Maximum data were obtained from Chelsa database (chelsa-climate.org). Latitude and longitude data were used for merging with the bioclimatic data. A resolution of 10-min was selected for this study. Bioclimatic variables (BIO1 – BIO19) were derived from monthly temperature and rainfall values representing annual trends, seasonality, and extreme environmental factors (Fick & Hijmans, 2017). The interpretation of bioclimatic variables is provided in Table S8. Future data were obtained similarly and will be clarified more in the “Species Distribution Modelling” section below.

### Regression Analysis

To determine which bioclimatic variables influenced the strength of secondary dormancy, data were analyzed using a generalized linear mixed model (GLMM) with a binomial distribution and individual relatedness as random effect. Secondary dormancy was modeled as a binomial response variable, defined by the number of germinated seeds (successes) relative to the total number of seeds per replicate. Thus, dormancy was analyzed as a binary outcome (success/failure) at the seed level using a logistic regression framework, with climatic variables included as predictors.

In such a logistic regression model (binomial family with logit link function), the probability of success πi for the outcome yi is defined as: logit(πi) = ln[πi/(1-πi)] The linear predictor including fixed effects terms: ni = βo + β1 * (treatment) + β2 * (bioclimatic variable) + β3 *(treatment)* (bioclimatic variable) + ui where ni is germination rate; βo is the intercept; β1, β2, β3 are coefficients for treatments, bioclimatic variables, and their interactions, respectively; ui ∼ N(0, σ ^2^ K) is the random effect, where K is the kinship matrix, accounting for population structure. The kinship-correlated random effect was thereby controlling for potential confounding effects on associations between phenotypes and explanatory variables due to shared ancestry.

The model was fitted with *relmatGlmer* function of package *lme4qtl* (Ziyatdinov et al., 2018) using logit link function. The kinship matrix had to be converted to positive definite matrix, using *nearPD* function of *Matrix* package (R Core Team, 2021). The resulting model coefficients reflect the effect size of each predictor on dormancy variation.

### Species Distribution Modelling

We used species distribution model (SDM) to compare past, present and future habitat suitability of two ecotypic groups split around the median of corrected secondary dormancy measured in trial 1 (i.e., the residuals of secondary dormancy regressed on primary dormancy for the trial where secondary dormancy was the strongest). The aim of the SDM analysis was to evaluate whether these two groups had equal likelihood to find suitable habitats over time. Genotypes of the strong-secondary-dormancy group had values above the trial 1 median, whereas genotypes of the weak-secondary-dormancy group had values below the trial 1 median. We used the R/biomod2 package (Thuiller et al., 2009) with the parameters explained in the following four steps. This procedure and the results are reported according to the standard protocol proposed by (Zurell et al., 2020).

Step 1 Overview: The spatial extent was limited from –10^0^ to 50^0^ for longitude and from 35^0^ to 70^0^ for latitude. The biodiversity data type was set on presence-only, with the locations of the samples marked as present. For predictor variables, we selected bioclimatic factors that are significantly associated with dormancy. The spatial resolution was set to 10-minute.

Step 2 Data preparation: As only presence data were available, pseudo-absence data needed to be created, because presence/absence data are required for most SDM algorithms. To ensure these pseudo-absences were environmentally distinct from presence locations, we used the Surface Range Envelope (SRE) strategy within the biomod2 framework. This method excludes areas with environmental conditions similar to those of the presence points, reducing the risk of misclassifying potential presences. We generated three sets of pseudo-absences, with the number of pseudo-absences equal to the number of presence points in each replicate, following the recommendations of Barbet-Massin et al. (2012).

For future data, we chose the representative concentration pathway (RCP) of 4.5 from the Coupled Model Intercomparison Project Phase 5 (CMIP5). RCP 4.5 corresponds to an intermediate scenario in which emissions peak at around 2040 and then decline. RCP 4.5 – the most probable scenario given that no climate policies are applied – considers non-renewable fuel availability. All bioclimatic variables were tested for multi-collinearity. We tested all predictor variables for multicollinearity and found that none were highly correlated (|r| < 0.7); thus, no variable re-selection or dimensionality reduction (e.g., PCA) was required.

Step 3 Model options: We tested different model algorithms, namely, Random Forest (RF), Gradient Boosting Machine (GBM), Generalized Linear Model (GLM), and Maxent. For calibration and evaluation, we implemented 3-fold cross-validation using a random splitting approach, whereby 80% of the data were used for training and 20% for testing in each replicate. This corresponds to random k-fold cross-validation, a standard method for evaluating model robustness when spatial autocorrelation is not the primary focus. We adopted this approach as our primary goal was to assess differences in habitat suitability between genotype groups. Assuming these two groups have similar dispersal abilities, the limitation of SDM applies equally to the two groups. We were aware of imperfect detection because we had sampling bias issues, that is, not all *A. thaliana* in the given geographical range were sampled. The sampling of individuals in the Central European region was less dense than that in Sweden and Spain. Cross-validation, which represents the random effects of selecting data, can show the sensitivity of the models to the input data. The evaluation metrics included TSS, ROC, and KAPPA scores.

Step 4 Assessment and Prediction: the biomod2 package examines the importance of each variable in the final model. Once the model is trained, a standard prediction is made, one variable is randomized, and a new prediction is made. The correlation score between the new prediction and standard prediction is given as an estimation of the variable importance in the model. The influence of each variable was visualized by plotting response curves. The ensemble modelling option combines individual models to build a metamodel. Models with TSS, ROC, and KAPPA lower than 0.8 were excluded. We used binary projection using ROC and a so-called weighted mean, which was done for all chosen models for each run, to project current and future climate models.

### Genome-wide Association Analysis

Variant calling was performed from raw sequence data for all 361 lines simultaneously. Genomic data of 309 genotypes from the 1001Genome data set are available on the European Nucleotide Archive (ENA), and data of 52 accessions were obtained from Wieters et al., 2021 (Suppl. Table S1 for accession information). The SRA files were downloaded from ENA using prefetch and converted into FASTQ using fastq-dump (from sra toolkit, US National Institutes of Health, 2023). Low-quality signals were detected, and polyG and sequencing adapters were removed using fastp (Chen et al., 2018). For this, the minimum read length was set to 50bp, the minimum Phred quality score was 15, and the quality threshold was 40%. The sequences were mapped to the reference genome (TAIR10, arabidopsis.org) using bwa-mem (Li, 2013) and converted to bam files using samtools (Danecek et al., 2021). The mapping quality was evaluated using multiqc (Ewels et al., 2016). Variant calling was performed using bcftools mpileup and bcftools call (Danecek et al., 2021; Li, 2011). The minimum mapping quality was set to 20, and the minimum base quality was set to 30. SNP data were processed with bcftools, vcftools, and PLINK for minor allele frequency filtering (0.05), maximum missingness (0.95), minimum depth and maximum depth (minDP = 10 and maxDP = 50), indels and low-quality calls (< 30), and prune linkage disequilibrium (r > 0.1, scanning 50kb window, 10bp step size) (Danecek et al., 2011; Purcell et al., 2007). Genome-wide association analysis (GWAS) was conducted to identify genetic variants associated with secondary dormancy. Association analysis was performed with GEMMA using PLINK files (Zhou and Stephens, 2012). Kinship matrix was calculated using KING algorithm of PLINK, and GWAS was performed using the Mixed Linear Model (MLM) approach (-lmm 4). The –log10 p-values of the model were adjusted using Bonferroni correction at a significance level of 5%. Chromosomal position and candidate genes were checked using the available annotated genes in the TAIR database (arabidopsis.org). We controlled for population structure using kinship matrix and added primary dormancy as a covariate to control for seed age difference across the trials. We combined the p-values from the different GWAS trials using Fisher’s combined probability test, which provides a robust approach for synthesizing results from multiple studies (Walsh and Lynch, 2018). Fisher’s method is based on the assumption that the p-values under the null hypothesis are uniformly distributed between 0 and 1. The method transforms each p-value using the natural logarithm and then sums these transformed values. It computes a combined test statistic χ^2^ given by

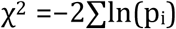

where p_i_ represents the p-values from the individual trials. This test statistic follows a chi-square distribution with 2k degrees of freedom, where k is the number of studies combined. We used Python and libraries namely numpy (Haris et al., 2020), scipy.stats (McKinney, 2010), and pandas (Virtanen et al., 2020).

## Results

### Secondary dormancy variation correlates with primary dormancy

To quantify genetic variation in heat-induced dormancy with latitude, we used seeds collected from 361 European accessions of *A. thaliana* grown in a common garden. Seed dormancy was measured in three trials, with seeds sampled from the same seed batch. The trials differed in the number of genotypes that were analyzed: 295, 361 and 344 genotypes, and seeds were six, twelve and twenty-four months after-ripened in trials 1, 2, and 3, respectively. We implemented three treatment groups for the germination tests: a germination test without any treatment to assess primary dormancy; a germination test following exposure to 4°C to evaluate seed viability, hereafter the control treatment; and a germination test after sequential exposure to 4°C and 37°C to assess secondary dormancy (Figure 1).

Both the number of lines exhibiting secondary dormancy and the strength of this dormancy decreased across the experiments (Figure 2). Seed viability remained high after 12 months of storage, because the control germination rate approached 100% after cold treatment. This rate, however, dropped to 50% for seeds after 24 months of seed storage. Primary dormancy was progressively reduced from Trial 1 (6-month-old seeds) to Trial 2 (12 months old seeds) and almost completely lost in Trial 3 (24-month-old seeds) (Figure 2).

**Figure 2.**
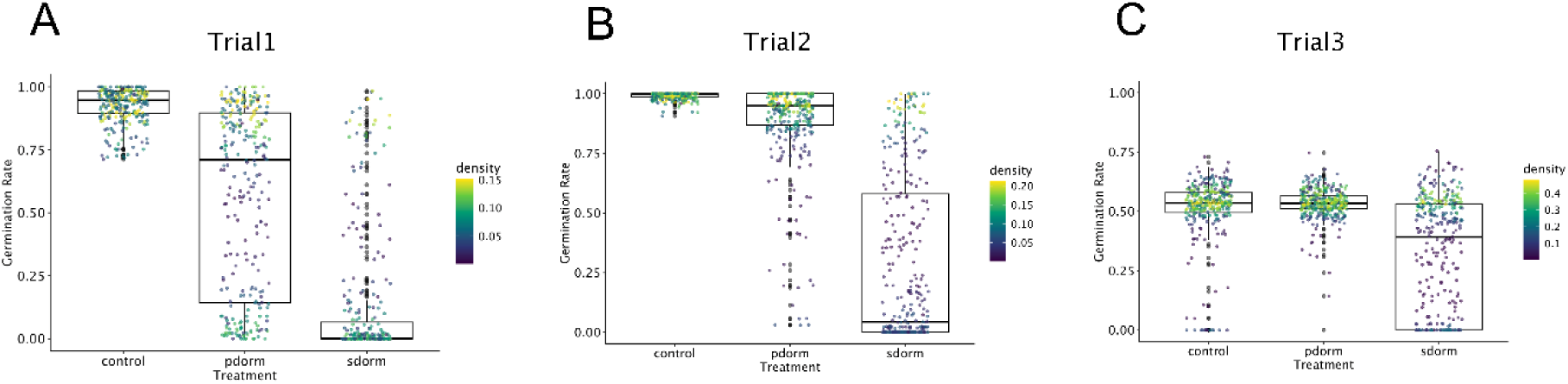
Germination rates across three treatments over three trials. Germination rates were measured for primary dormancy (pdorm), secondary dormancy (sdorm), and control treatments across three trials. The color gradient represents the density of data points. (A) Trial 1: conducted in May 2021 (6 months post-harvest). (B) Trial 2: conducted in December 2021. (C) Trial 3: conducted in December 2022. The sdorm” treatment represents germination rates following a 3-day stratification at 4°C to release dormancy, followed by a 4-day treatment at 37°C. The pdorm treatment tested germination rates without any pre-treatment, whereas the control treatment involved seeds stratified for 3 days at 4°C to release dormancy before testing germination. All germination tests were conducted under long-day conditions at 20°C, and germination rates were recorded after 7 days.

Levels of seed secondary dormancy were consistently correlated across trials (p < 2.814e-09, Suppl. Table S2). While primary dormancy was released by stratification prior to heat exposure, secondary dormancy remained significantly correlated with primary dormancy in all three trials (maximum p = 0.00135, Suppl. Table S3). However, the strength of this correlation declined over time (Trial 1: Spearman rho = 0.427; Trial 3: Spearman rho = 0.172). Residual variation also increased across trials, confirming the increasing divergence from primary dormancy signal (Suppl. Figure S2B).

### Bioclimatic drivers of heat-induced secondary dormancy

Despite the variation in levels of dormancy observed across trials, secondary dormancy decreased with increasing latitude in all three trials (Figure 3, Suppl. Table S4). This correlation may arise from the history of post-glaciation expansion in the species, from the indirect correlation to primary dormancy, or from the specific adaptation of secondary dormancy along climatic clines. To discriminate between these hypotheses, we tested whether the geographic distribution of secondary dormancy variation was significantly associated with climatic variables after accounting for individual relatedness and both variance in primary dormancy and variance in germination rate. Using publicly available global climate data with 10-minute resolution in longitudinal and latitudinal coordinates, we conducted a generalized linear mixed model (GLMM) analysis to investigate the association between 19 bioclimatic variables and germination outcome (Figure 4).

**Figure 3.**
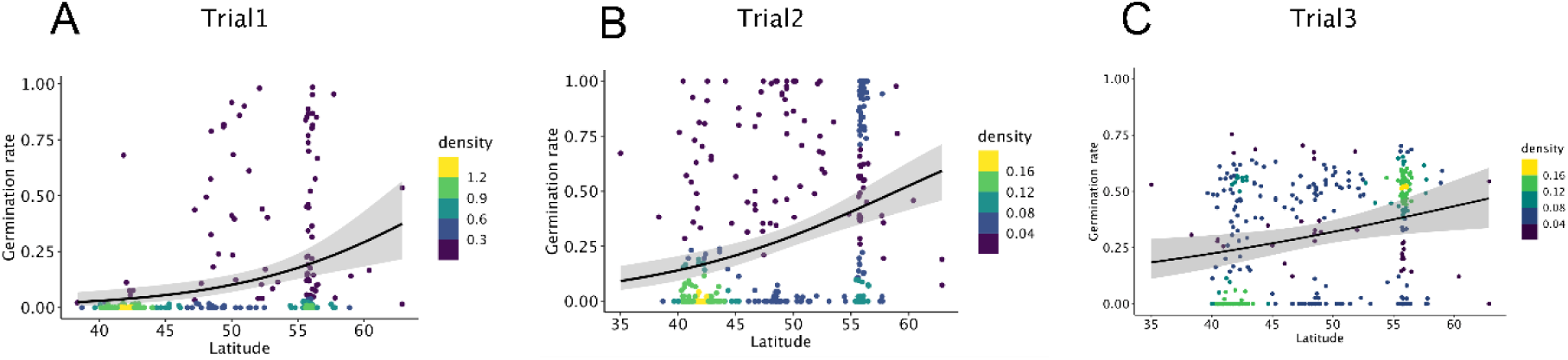
Germination rates of heat-induced secondary dormancy treatment as a function of latitude of origin. The secondary dormancy treatment represents germination rates following a 3-day stratification at 4°C to release dormancy, followed by a 4-day treatment at 37°C. Germination tests were conducted under long-day conditions at 20°C, and germination rates were recorded after 7 days. The correlation between germination rate and latitude of origin was assessed for three trials: (A) Trial 1, (B) Trial 2, and (C) Trial 3. Original values were used without any corrections or adjustments. Spearman’s rank correlation coefficients (rho) and corresponding p-values were as follows: (A) rho = 0.386, p = 6.823 × 10^-12^; (B) rho = 0.339, p = 2.986 × 10^-11^; and (C) rho = 0.278, p = 1.144 × 10^-7^. Color density highlights regions with increasingly high concentration of values.

**Figure 4.**
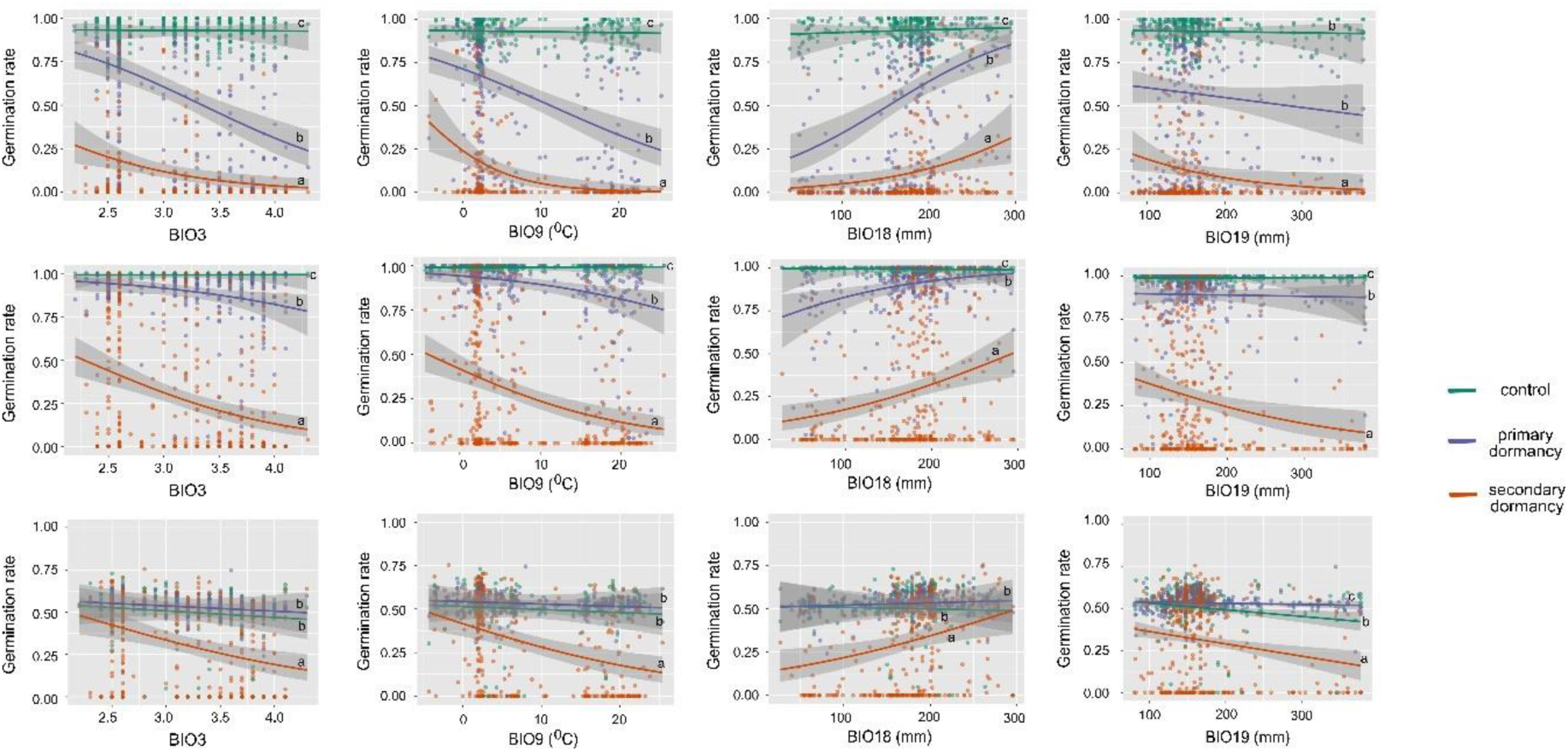
Germination rate across three treatments correlated with four bioclimatic variables (BIO3, BIO9, BIO18, and BIO19). The relationship between germination rates and four bioclimatic predictors—BIO3 (isothermality), BIO9 (mean temperature of the driest quarter), BIO18 (mean precipitation of the warmest quarter), and BIO19 (mean precipitation of the coldest quarter)—across three trials: (A) Trial 1, (B) Trial 2, and (C) Trial 3. The so-called corrected secondary dormancy treatment represents germination rates following a 3-day stratification at 4°C to release dormancy, followed by a 4-day treatment at 37°C, corrected for the residual effects of primary dormancy. The primary dormancy treatment tested germination rates without any pre-treatment, and the control treatment involved seeds stratified for 3 days at 4°C to release dormancy before testing germination. All germination tests were conducted under long-day conditions at 20°C, and germination rates were recorded after 7 days. The bioclimatic variables include BIO3, which is unitless; BIO9, measured in degrees Celsius; and BIO18 and BIO19, measured in millimeters.

We found that genotypes exhibiting strong secondary dormancy tended to originate from locations associated with increasing mean temperatures and decreasing rainfall (mean temperature of driest quarter – BIO9 values). Summer precipitation (mean precipitation of warmest quarter – BIO18 values) was negatively correlated with secondary dormancy (mimimum p < 2.2e-16, Suppl. Table S5). Additionally, we observed a positive correlation between winter precipitation (mean precipitation of coldest quarter – BIO19 values) and heat-induced secondary dormancy (maximum p = 3.55e-8, Suppl. Table S5). Variation in secondary dormancy is further associated with climatic fluctuations. We observed that higher isothermality (BIO3)—indicative of greater diurnal temperature variation relative to annual temperature range—was significantly associated with increased levels of secondary dormancy (maximum *p* = 1.77e-6; Suppl. Table S5). Altogether, these results suggest that *A. thaliana* populations evolved stronger heat-induced secondary dormancy in environments characterized by low precipitation, high temperatures, and pronounced short-term temperature fluctuations, conditions likely serving as ecological cues signaling germination stress (Figure 4, Suppl. Figure S3, Suppl. Table S5). Since this correlation of phenotypic variation with environmental parameters was significant after accounting for population structure (see Methods), we conclude that the distribution of genetic variation for heat-induced secondary dormancy was shaped by natural selection.

Primary dormancy variation was also associated with isothermality (BIO3), mean temperature of driest quarter (BIO9), and summer precipitation (BIO18) in the first two trials (maximum p < 2.2e-16, Suppl. Table S5), but not with winter precipitations (BIO19) (maximum p = 0.16196, Suppl. Table S5). In the third trial, only BIO19 was associated with remaining primary dormancy variation (p = 0.00143, Suppl. Table S5). As primary dormancy decreased across trials, the slope of its correlation with bioclimatic variables also decreased (Suppl. Table S5, Figure 4). Interestingly, the slope of the relationship between secondary dormancy and each of these four climatic variables was always significantly stronger than the relationship with primary dormancy, irrespective of the level of residual primary dormancy (Figure 4). The shared association between secondary dormancy and primary dormancy with temperature– and rainfall-related bioclimatic variables suggests that these traits may constitute an adaptive dormancy syndrome that coordinates dormancy responses to seasonal and climatic fluctuations. The sharper climatic clines of secondary dormancy compared to primary dormancy suggest that secondary dormancy is a better descriptor of local adaptation than primary dormancy.

### Relative size of suitable habitats is predicted to increase for high secondary dormancy ecotypes

Since secondary dormancy levels displayed a pattern of local adaptation along several bioclimatic gradients, we reasoned that the fitness of high vs. low secondary dormancy ecotypes would depend on the relative size of their suitable habitat (Banta et al. 2012, Ikeda et al. 2016). Using the four identified bioclimatic factors shown above to drive local adaptation of secondary seed dormancy (BIO3, BIO9, BIO18, BIO19), we tested quantified the amount of suitable habitat inferred by projecting species distribution models for both ecotypic groups (Figure 5). We used current data to determine the relationship of each ecotype with bio-climatic variables drawing their respective niches. We then used past climate data from the Last Glacial Maximum and future climate data based on RCP4.5 (see Methods).

**Figure 5.**
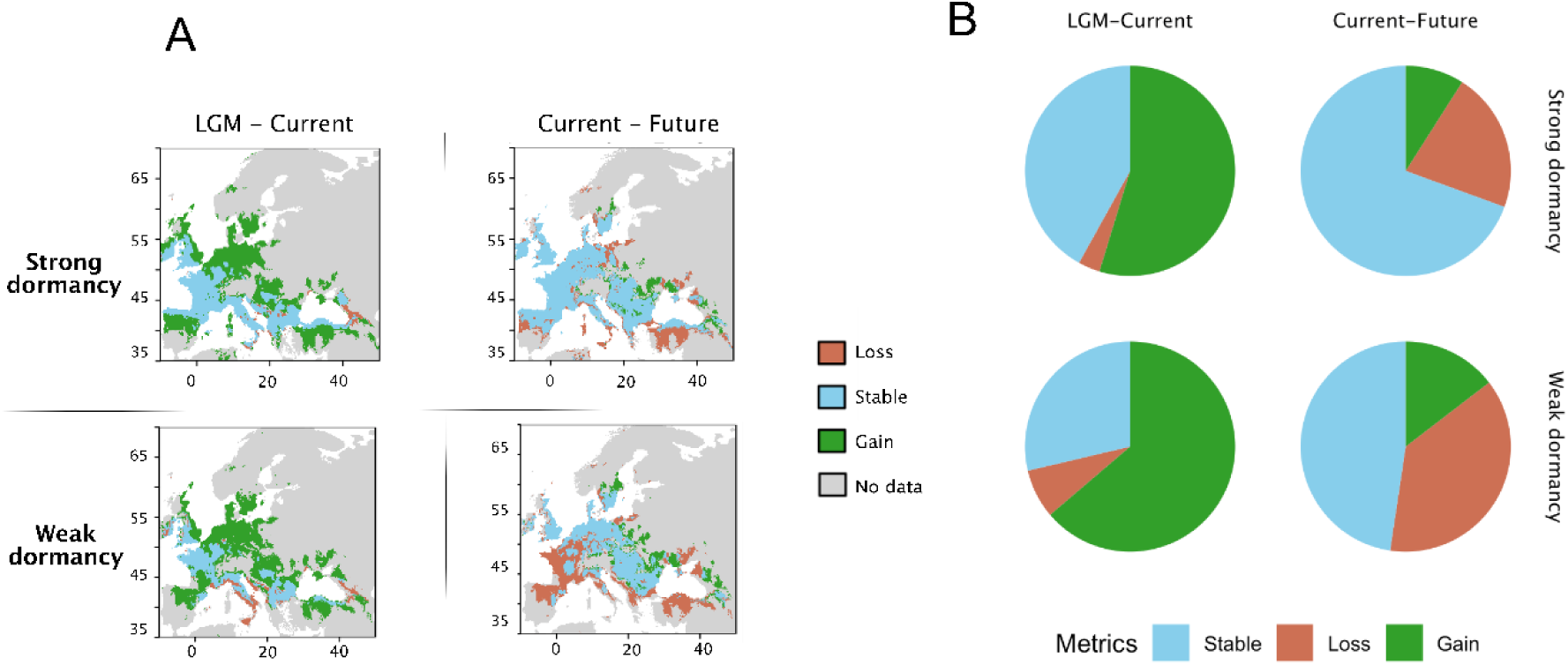
Species distribution model (SDM) for strong and weak heat-induced secondary dormancy genotypes. The environmental envelopes and projected range changes for genotypes with strong and weak secondary dormancy across three time periods: the Last Glacial Maximum (LGM), the present, and the year 2050 under projected climate scenarios. To construct the SDM, four bioclimatic variables were used: temperature seasonality (BIO3), mean temperature of the driest quarter (BIO9), precipitation of the warmest quarter (BIO18), and precipitation of the coldest quarter (BIO19). Data from Trial 2, the most complete dataset, was used for the analysis. Genotypes were divided into strong and weak secondary dormancy groups based the secondary dormancy corrected for primary dormancy, so-called corrected secondary dormancy, which represents germination rates following a 3-day stratification at 4°C to release dormancy, followed by a 4-day treatment at 37°C, corrected for the residual effects of primary dormancy. The figure presents species range changes in three categories—loss (orange), stable (blue), and gain (green)—alongside areas (gray) that have no data due to insufficient number of data points in those regions (e.g., the northeastern Europe). Results are derived from an ensemble model, which combines the best-performing model based on ROC metrics and calculates weighted means across three cross-validation runs. (A) Species range changes for strong secondary dormancy genotypes (upper) and weak secondary dormancy genotypes (lower) during two transitions: LGM to the present (left) and the present to 2050 (right). (B) Pie charts depicting the proportion of habitat loss, stability, and gain for each group during these transitions. Statistical analyses reveal significant differences in range change patterns between strong and weak dormancy genotypes (chi-squared test: LGM-current: X² = 200.66, df = 2, p < 2.2 × 10^-16^; current-future: X² = 4484, df = 2, p < 2.2 × 10^-16^).

Among the different models we tested, the best model performance was obtained with Random Forest (ROC and TSS scores > 0.95; Suppl. Figure S4). As anticipated, given its strong influence on dormancy ecotypes, BIO9 determined about 40% of the distribution of both strong– and weak-secondary dormancy ecotypes (Suppl. Figure S5). BIO9 is particularly influential because seeds experience both heat and drought during this period, making the triggering of dormancy-related responses crucial for seedling survival. Furthermore, BIO9 likely interacts with other predictors, such as precipitation variables (e.g., BIO18 and BIO19), shaping the seasonal microclimatic conditions that define the ecological niches in which dormancy strategies confer an advantage.

Our model showed both the proportion of stable habitats for both ecotypic groups from LGM to current time, as well as the change in habitat boundaries. Past and predicted habitat range change depended on the secondary seed dormancy strategy (Figure 5B, LGM-Current: X-squared = 108.03, df = 2, p-value < 2.2e-16; Current-2050: X-squared = 814.45, df = 2, p-value < 2.2e-16). Habitat suitability was quantified by the amount of habitat, measured as pixel in the grid, that fits the current distribution of the two secondary dormancy types. Habitat suitability increased for strong dormancy ecotypes in northwestern and southern areas of Europe such as in the Balkans and Spain, compared to weak secondary dormancy (measured in pixels). In total, weak secondary dormancy genotypes appeared to have benefitted from a larger habitat gain due to significant gains in central and northeastern Europe. However, in the close future, only strong dormancy ecotypes are predicted to maintain most of their current habitat, weak secondary dormancy ecotypes, instead, are predicted to lose around half of their current habitat (primarily in western and southern Europe) with only slight gains (in northeastern Europe) (Figure 5B, LGM – current: X-squared = 200.6, df = 2, p-value < 2.2e-16, current – 2050: X-squared = 4484, df = 2, p-value < 2.2e-16). In summary, the climate projection, which here is based on an intermediate scenario with emission peak around 2040, indicates that ecotypes with strong secondary dormancy should lose less habitat than those with weak secondary dormancy. Strong secondary dormancy ecotypes are therefore predicted to be more resilient to expected environmental changes. Based on the size of predicted suitable habitats in changing climatic conditions, high-secondary dormancy ecotypes are predicted to make an ever increasing fraction of the *A. thaliana* population. Assuming that high– and low-dormancy ecotypes do not differ in their ability to migrate to new habitats, these findings suggest that high-dormancy genotypes have a fitness advantage.

### Genetic variants associated with the environment and secondary dormancy

To investigate the genetic basis of the dormancy variation revealed in our experiment, we performed genome-wide association studies (Figure 6). For primary dormancy, we observed a significant peak located only 14bp away from the region of the DELAY OF GERMINATION 1 (DOG1) gene, which was not found to associate with secondary dormancy (Suppl. Figure S6D; Suppl. Table S6). In order to identify the specific genetic basis of secondary dormancy, we accounted for variation in primary dormancy in the model of genetic association. We identified several SNPs with significant associations, notably, on chromosomes 2, 3, and 4 (Suppl. Figure S7, Suppl. Table S7), as annotated in Figure 6A. The location of these three associated SNPs did not overlap with the genomic regions associated with primary dormancy (Figure 6B). SNPs on chromosomes 2 and 4 did not fall close to any obvious candidate gene for secondary dormancy, but they may represent novel loci or the regulatory elements involved in dormancy-related pathways that have yet to be characterized.

**Figure 6.**
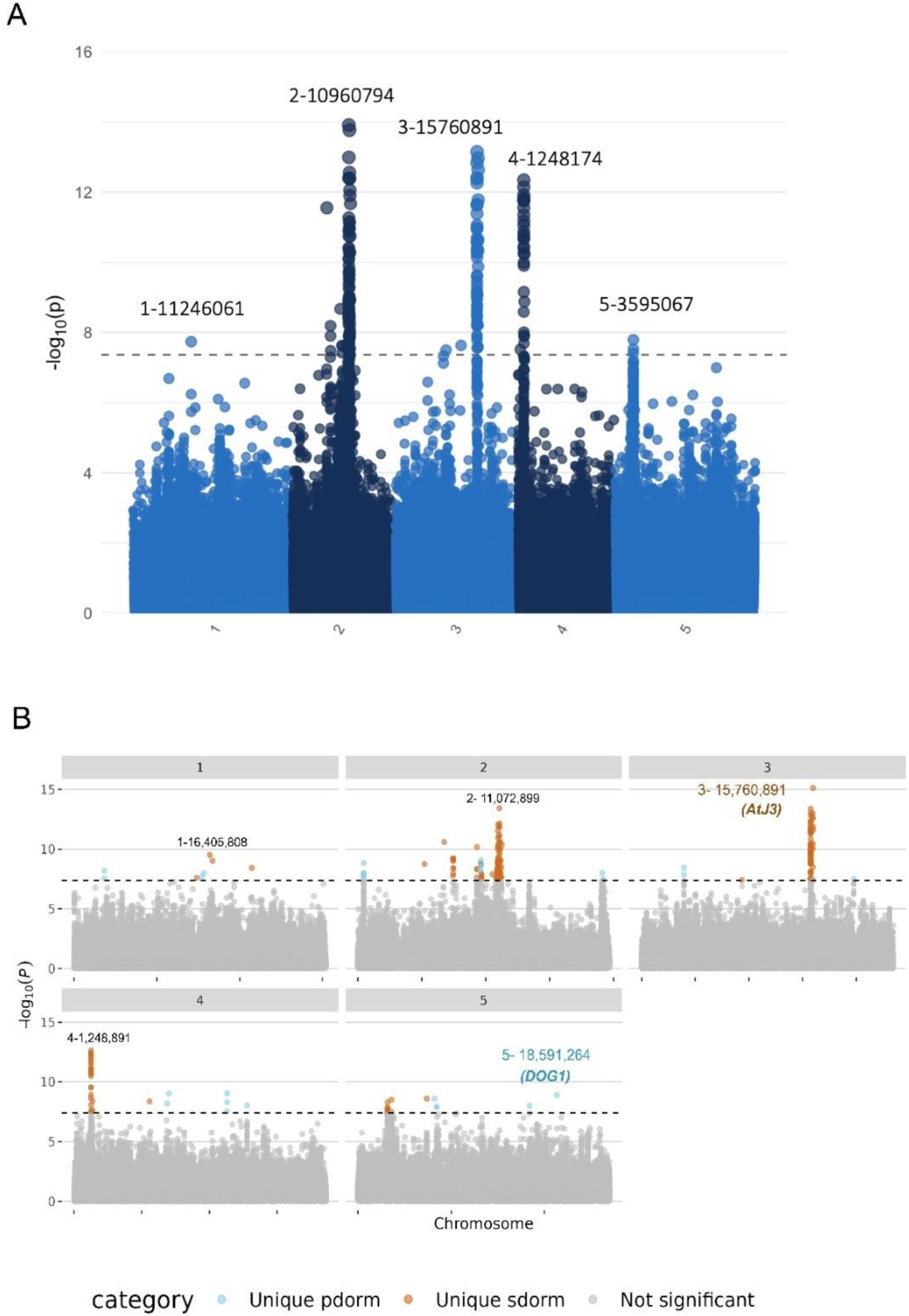
GWAS results for heat-induced secondary dormancy and primary dormancy SNP associations. (A) Manhattan plot illustrating the association of 1.2 million SNP markers with heat-induced secondary dormancy, using primary dormancy as a covariate. In the primary dormancy treatment, seeds were tested directly without any pre-treatment to assess the baseline dormancy established during seed maturation. For the secondary dormancy treatment, seeds underwent stratification at 4°C for three days to release dormancy, followed by a heat stress of 37°C for four days to induce secondary dormancy. The results are based on a combined p-value GWAS analysis of three trials, calculated using Fisher’s combined probability formula. The dashed horizontal line represents the significance threshold after applying a 5% Bonferroni correction. Significant SNPs crossing this threshold highlight genomic regions associated with secondary dormancy independent of primary dormancy. (B) Overlay of two Manhattan plots to compare SNP associations for heat-induced secondary dormancy without covariates (orange) and primary dormancy (blue). SNPs uniquely associated with primary dormancy are highlighted in blue (unique pdorm), and SNPs uniquely associated with secondary dormancy are highlighted in orange (unique sdorm). Gray dots represent SNPs that do not meet the 5% Bonferroni correction significance threshold for either dormancy trait.

The SNPs on chromosome 3 were particularly intriguing, because they were located approximately 100kb upstream and 60kb downstream of the DNAJ Homolog 3 gene, (AT3G44110, AtJ3, or AtDjA3). Although several genes are present in this region, AtJ3 encodes for a heat-shock co-chaperone that is involved in the control of germination in stressful conditions, making it a likely candidate (Salas-Munoz et al., 2016; Barghetti et al., 2017; Wu et al., 2019). Several elements indicate that the absence of associated SNPs within and or close to the gene may be due to mapping issues. The gene is surrounded by numerous repeats that perturb the quality of mapping. In addition, in the Col-0 reference genome, substantial deletions in the gene further complicate SNP detection within the gene body.

## Discussion

In this study, we investigated variation in heat-induced secondary dormancy in a European-wide collection of *A. thaliana* genotypes. Germination timing is modulated by seed dormancy, which allows seeds to avoid unfavorable conditions (Baskin and Baskin, 1998). Primary dormancy, formed during seed maturation, is well-studied and contributes to adaptation to drier climates (Kronholm et al. 2012; Chiang et al., 2013; Debieu et al., 2013; Kerdaffrec and Nordborg, 2017). Here, we shed new light on the genetics of heat-induced secondary dormancy, its covariation with primary dormancy, as well as its relevance in adaptation to climate.

Our study confirms previous findings, namely, that heat-induced secondary dormancy depends on the degree of primary dormancy formed in the mother plant (Auge et al., 2015). Our genetic analysis highlights the existence of both common and specific pathways controling secondary dormancy. Here, primary dormancy was released by stratification at 4°C before heat exposure. Despite full primary dormancy release, variation in secondary dormancy still depended on the variation in primary dormancy of unstratified seeds. This relationship was strong when secondary dormancy was assessed in seeds with high primary dormancy and persisted in after-ripened seeds, although heat-induced secondary dormancy levels were lower. Heat-induced secondary dormancy thus appears to share the molecular mechanism that activate primary dormancy, a hypothesis that is in line with the study by Coughlan et al. (2017). However, GWAS controlling for primary dormancy identified a genomic region around *AtJ3* loci that is associated with secondary dormancy induced after seed stratification, indicating that additional molecular components distinct from those controlling primary dormancy also shape variation in secondary seed dormancy. As mentioned above, the *AtJ3* (AT3G44110) gene encodes a heat-shock chaperone involved in flowering time and germination (Salas-Munoz et al., 2016; Barghetti et al., 2017; Wu et al., 2019). In particular, the germination of *AtJ3* mutants is impaired in stress and effectively blocked by ABA treatment, two observations that support its possible role in secondary dormancy (Salas-Munoz et al., 2016). Intriguingly, its association with secondary dormancy suggests that pathways involved in temperature sensing and stress responses may play a role in regulating the seed’s ability to enter dormancy under unfavorable conditions. In addition, the variation in primary dormancy associated with variants close to *DOG1*, a well-known QTL for primary dormancy (Bentsink et al., 2006). *DOG1* was shown to explain much of the primary dormancy variation in Europe and Scandinavia (Kronholm et al., 2012; Kerdaffrec and Nordborg, 2017). Yet, associations with *DOG1* are sometimes elusive. For example, it was not associated with primary dormancy but with cold-induced secondary dormancy and flowering time, in a study of 559 *A. thaliana* genotypes (Martinez-Bedeja et al. 2020). Since here, we found that *DOG1* variants were associated with primary dormancy but not with heat-induced secondary dormancy, it appears that *DOG1* is not responsible for the covariation between the two traits, as measured in this study. Further work will be needed to elucidate the molecular basis of secondary dormancy and clarify the role of *DOG1* variation. In this effort, our data clearly show that the dependency of secondary dormancy levels on the degree of seed after-ripening will have to be carefully accounted for.

The significant covariation of secondary dormancy with climatic variables, such as summer heat and precipitation, further affirms the adaptive relevance of secondary dormancy. *Arabidopsis thaliana’s* range expansion from southern and eastern European refugia to northern areas has created a gradient of genetic variation across the continent (Fulgione & Hancock, 2018; 1001 Genomes Consortium, 2016). By controlling for this gradient, our results demonstrate the adaptation of secondary dormancy to local climates, a result that contrasts with a previous study by Ibarra et al. (2016) that did not account for population structure. Heat thus induced higher levels of secondary dormancy in genotypes originating from warmer climates with more frequent drought (Buijs et al., 2020; Postma et al., 2015). Indeed, in regions with harsh dry areas, a delay in germination was reported to increase seedling survival (Springthorpe and Penfield, 2015). Furthermore, we found associations of both primary and secondary dormancy with climatic variables that quantify the harshness of the summer season: isothermality, summer precipitation, and mean precipitation of the warmest quarter. Many studies also previously emphasized both the role of high temperatures in dormancy regulation and the effect of elevated temperature and disrupted precipitation on plant traits (Footitt et al., 2011; Hamann et al., 2021; Ibarra et al., 2016; Schmitz et al., 2023; Kerdaffrec and Nordborg, 2017; Kronholm et al., 2012; Postma et al., 2015; Zacchello et al., 2020, 2022). Microgeographic variation in temperature in urban habitats indicates that secondary dormancy levels correlate with temperature or environmental disturbance (Schmitz et al., 2023). Overall, covariation between primary dormancy, summer harshness, and seed age demonstrates that secondary dormancy adaptation involves diverse responses to complex environments.

Ecological specialization requires the optimization of trait suites, which are sometimes described as trait syndromes (Diaz et al., 2016; Takou et al., 2018). For example, a study in tropical and temperate ecosystems showed that plant species converge on similar defense syndromes based on shared environmental pressures while maintaining divergent strategies across syndromes to promote diversity (Kursar and Coley, 2003). Likewise, another study in Populus fremontii suggested a highly coordinated adaptation across multiple trait spectra to local climate constraints (Blasini et al., 2020). Trait syndromes can also vary within species (Takou et al., 2018). Primary dormancy has been shown to evolve as a trait syndrome along with growth rate and flowering time (Debieu et al., 2013; Takou et al., 2018). Here, two elements suggest that secondary dormancy forms an adaptive trait syndrome with primary dormancy. First, both dormancies are correlated, but GWAS reveals the distinct genetic bases of their variation. Second, they are both shaped by similar climatic variables, but the stronger pattern of association found for secondary dormancy suggests that the latter is crucial to improve the adaptation of germination strategy to local conditions.

In order to evaluate the relative fitness advantage provided by strong secondary dormancy, we used species distribution models to contrast the amount of past, current and future suitable habitats. Species distribution models will not accurately predict the distribution that will be achieved in past or projected climate because they ignore the potential of species to effectively colonize new habitats or to adapt genetically (Anderson et al. 2025, Nogues-Bravo, 2009, Fitzpatrick and Hargrove, 2009). The analysis conducted here illustrate well why this is the case. Indeed, the potential habitat suitability of locally adapted genotypes will differ (Ikeda et al. 2016, Banta et al. 2012). At the same time, our findings lend support to the notion that strong secondary dormancy will be more often advantageous than weak secondary dormancy. The bioclimatic variables associated with secondary dormancy are predicted to change rapidly in the near future. Selection driven by heat and drought will likely increase, especially in southern Europe (Exposito-Alonso et al., 2019). This study thus complements previous studies showing that the timing of germination is under strong selection and key to species distribution (Donohue et al., 2010; Huang et al., 2010; Willis et al., 2014, Exposito-Alonso, 2020) and highlights the role heat-induced secondary dormancy ecotypes may play in the response to climate change. Yet, an experimental validation of the adaptive relevance of secondary dormancy remains warranted. Testing the relative fitness advantage of genotypes with or without the ability to trigger secondary dormancy, however, would be challenging, because the trait requires tightly controlled conditions to be quantified although it is an adaptation to summer harshness, which describes an increased probability to face unsuitable conditions.

## Conclusions

This study describes the genetic variation of heat-induced secondary dormancy in *A. thaliana* and its ecological relevance in populations exposed to harsh summers. Secondary dormancy is influenced by primary dormancy but also involves distinct genetic components. Covariation with climatic variables, particularly temperature and precipitation, underscores the adaptive relevance of heat-induced secondary dormancy to local and future environments. Our findings suggest secondary dormancy is part of a broader dormancy trait syndrome, critical for aligning germination timing with environmental conditions experienced before and after dispersal. Understanding these dynamics is essential for predicting plant adaptation to future climates.

## Acknowledgments

We express special thanks to Cansu Aslan, for technical assistance, Lina Abdelwahed for discussions on species distribution modeling, Dr. Leen Nanchira Abraham for advice on bioinformatic workflows, and to Emily Wheeler for editorial assistance. This work was supported by the Deutsche Forschungsgemeinschaft (DFG, German Research Foundation) under Germany’s Excellence Strategy (EXC-2048/1-Project ID 390686111) and the research consortium TRR341.

## Author contributions

NLTT performed the experiments and data analysis; TA supervised species distribution modeling; GS assisted in experimental design and data presentation; JM supervised experiments, data analysis, and interpretation. NLTT, TA and JM jointly wrote the manuscript with feedback from all coauthors.

## Data Availability Statement

Raw data of germination tests are available on Figshare at https://doi.org/10.6084/m9.figshare.28218308.v1. Supplementary tables that were noted with “Attached as a separate file” are available on Figshare at https://doi.org/10.6084/m9.figshare.28409279.v1

Scripts used to carry out the analyses for this work are available at a Github repository (https://github.com/tntranloc/Athaliana_seed_secondary_dormancy).

## Benefit-Sharing Statement

Benefits from this research accrue from the sharing of our data and results on public databases as described above.

## Figures

**Figure S1.**
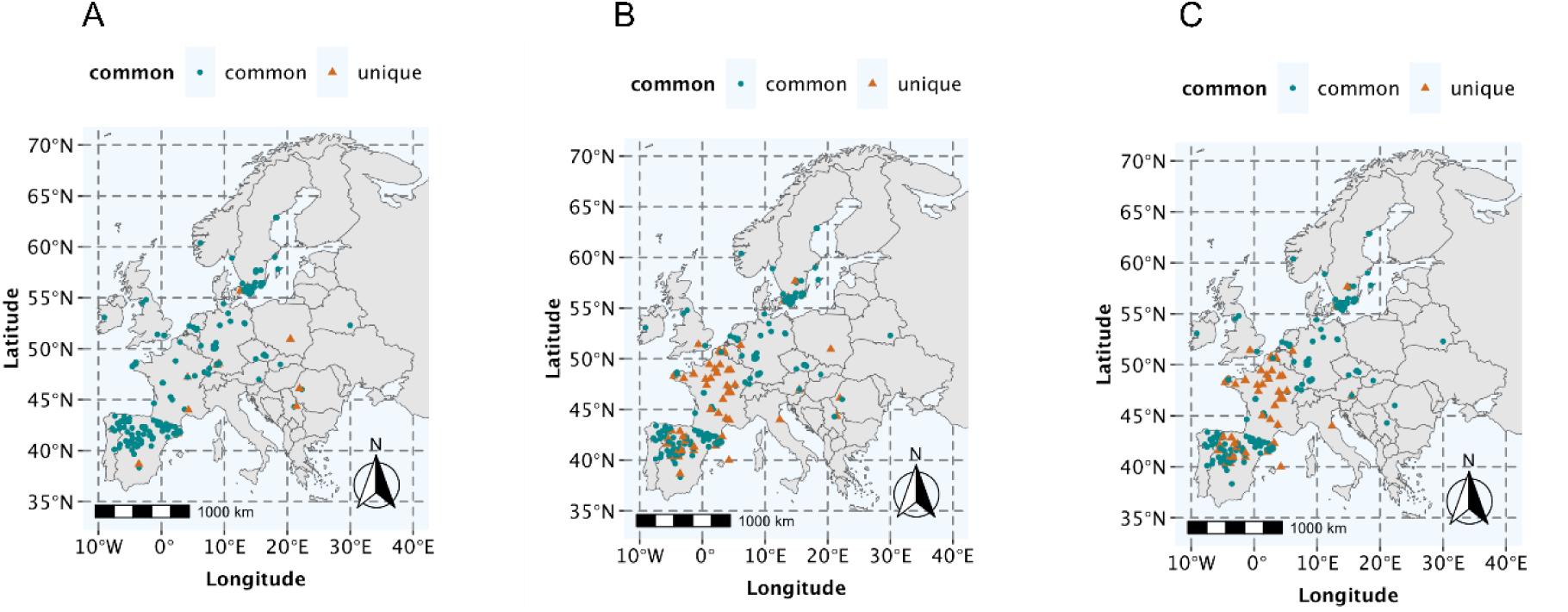
Geographic origin of European Arabidopsis thaliana accessions used in this study. The map displays the geographic locations of Arabidopsis thaliana accessions included in the study, shown separately for each trial. (A) Trial 1, (B) Trial 2, and (C) Trial 3. Green round markers represent common genotypes that are present across all three trials, while orange triangular markers indicate unique genotypes that are specific to a single trial.

**Figure S2.**
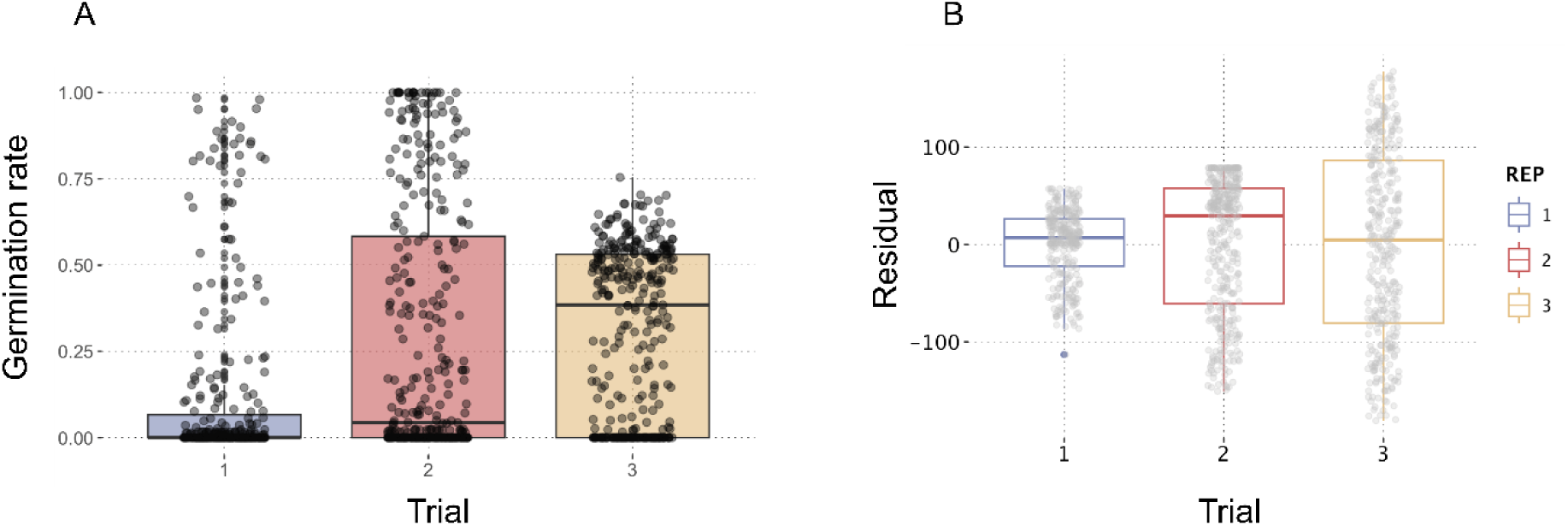
Temporal dynamics of heat-induced secondary dormancy. (A) Distribution of germination rates over time in response to the heat-induced secondary dormancy treatment across three trials. The x-axis represents time points corresponding to the three trials. Secondary dormancy treatment represents germination rates following a 3-day stratification at 4°C to release dormancy, followed by a 4-day treatment at 37°C. (B) Residuals of the linear regression of secondary dormancy rank on primary dormancy across the three trials. Primary dormancy tested germination rates without any pre-treatment. The residuals represent the component of secondary dormancy not explained by primary dormancy, highlighting the change in dormancy traits over time. All germination tests were conducted under long-day conditions at 20°C, and germination rates were recorded after 7 days.

**Figure S3.**
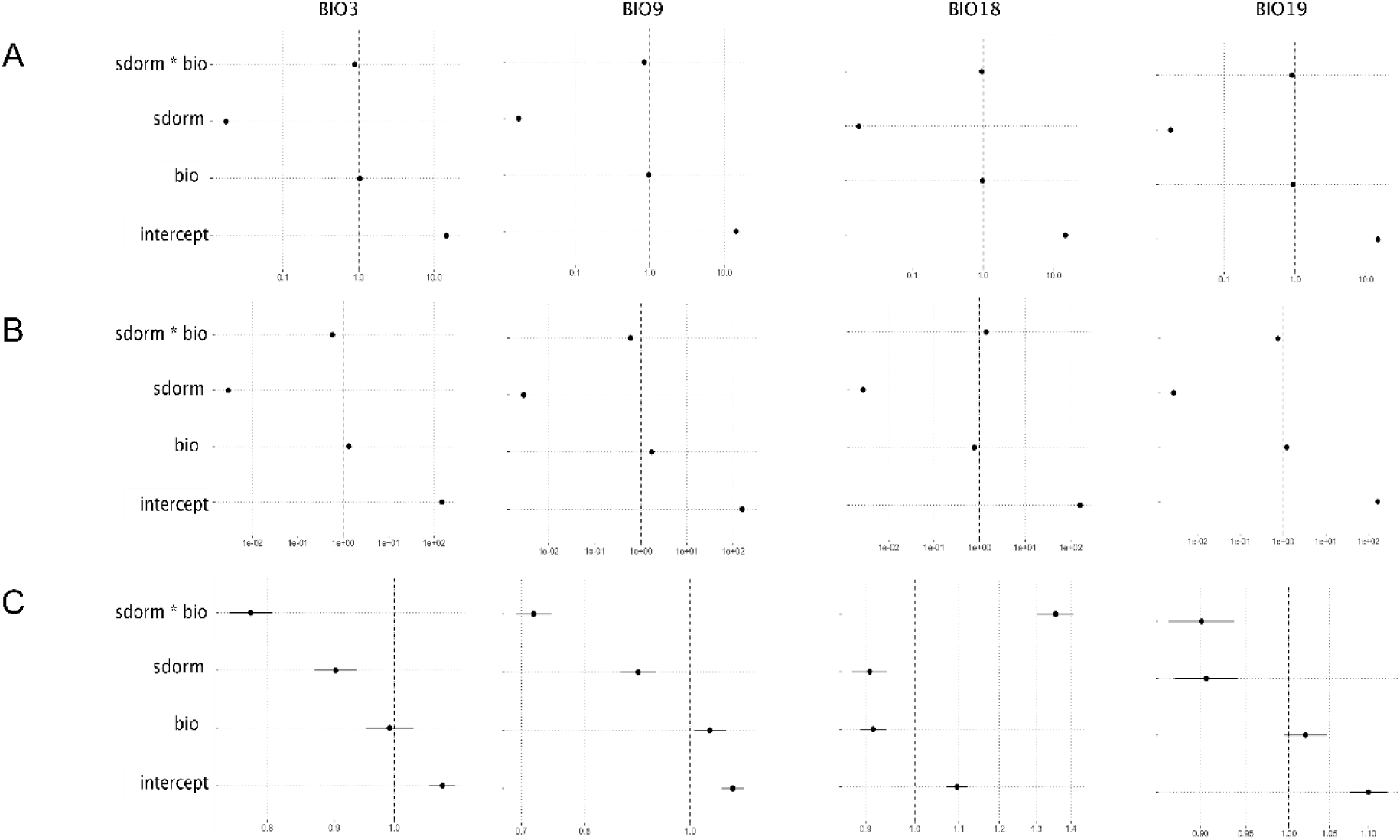
Bioclimatic variables as predictors of heat-induced secondary dormancy. Odds ratios are derived from binomial models with a logit link function for the effect of four bioclimatic variables—BIO3 (isothermality), BIO9 (mean temperature of the driest quarter), BIO18 (mean precipitation of the warmest quarter), and BIO19 (mean precipitation of the coldest quarter)—as predictors of secondary dormancy. Results are displayed separately for three replicates: (A) Trial 1, (B) Trial 2, and (C) Trial 3. Secondary dormancy (sdorm) was induced by a 4-day treatment at 37°C following a 3-day stratification at 4°C to release dormancy. Primary dormancy (pdorm) seeds were tested for germination without pre-treatment, and control seeds, used here as the reference level, were stratified for 3 days at 4°C and tested for germination. The sdorm values used in the models represent secondary dormancy corrected for primary dormancy (residuals of the regression of secondary dormancy on primary dormancy). The models assess the effect of each bioclimatic variable (bio) on germination outcomes under long-day conditions at 20°C, with germination tested after 7 days. BIO3 = isothermality (ratio; unitless); BIO9 = mean temperature of the driest quarter (°C); BIO18 = mean precipitation of the warmest quarter (mm); BIO19 = mean precipitation of the coldest quarter (mm).

**Figure S4.**
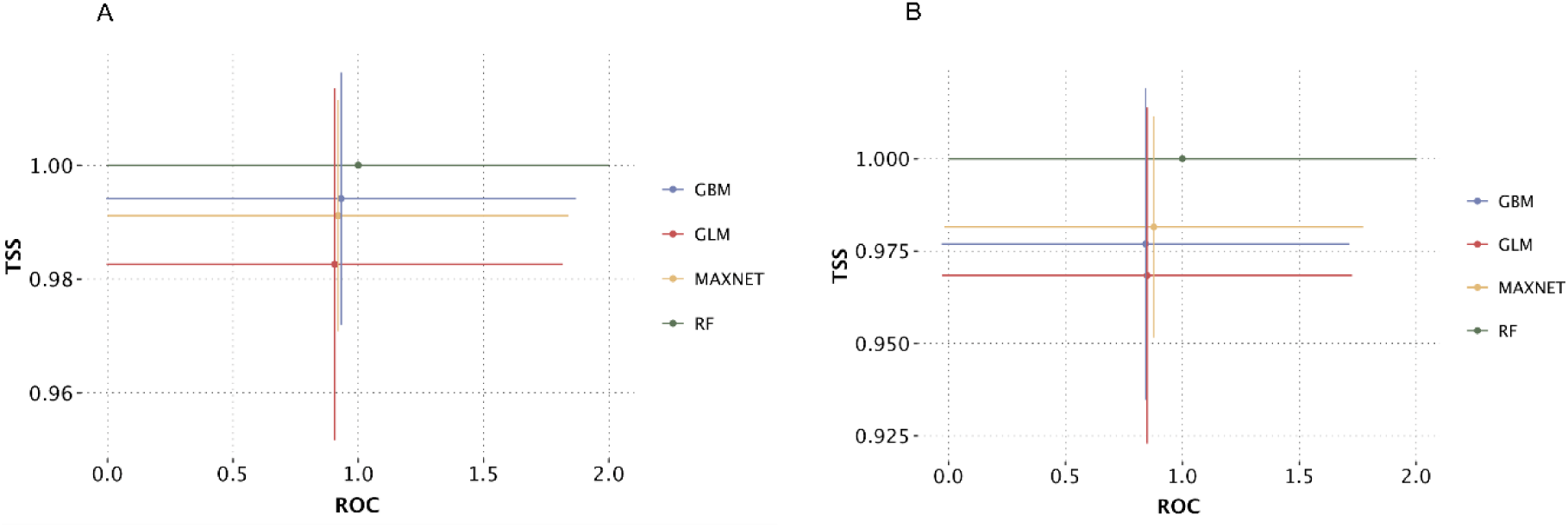
Performance comparison of modelling approaches using ROC and TSS metrics in predicting heat-induced secondary dormancy ecological niche. Comparisons of the performance of four modeling approaches: gradient boosting machine (GBM, blue), generalized linear model (GLM, red), maxnet (MAXNET, yellow), and random forest (RF, green). The comparison is based on two evaluation metrics: Receiver Operating Characteristic (ROC) and True Skill Statistic (TSS). The ROC metric assesses the model’s ability to distinguish between classes, according to which a higher value indicates improved ability to discriminate. TSS evaluates the model’s predictive skill, balancing sensitivity and specificity. These metrics provide a comprehensive evaluation of model performance across different approaches, facilitating the identification of the best method for predicting secondary dormancy. The color coding corresponds to the modeling approaches: blue for GBM, red for GLM, yellow for MAXNET, and green for RF. Each point on the plot represents the performance of a model under the respective metric.

**Figure S5.**
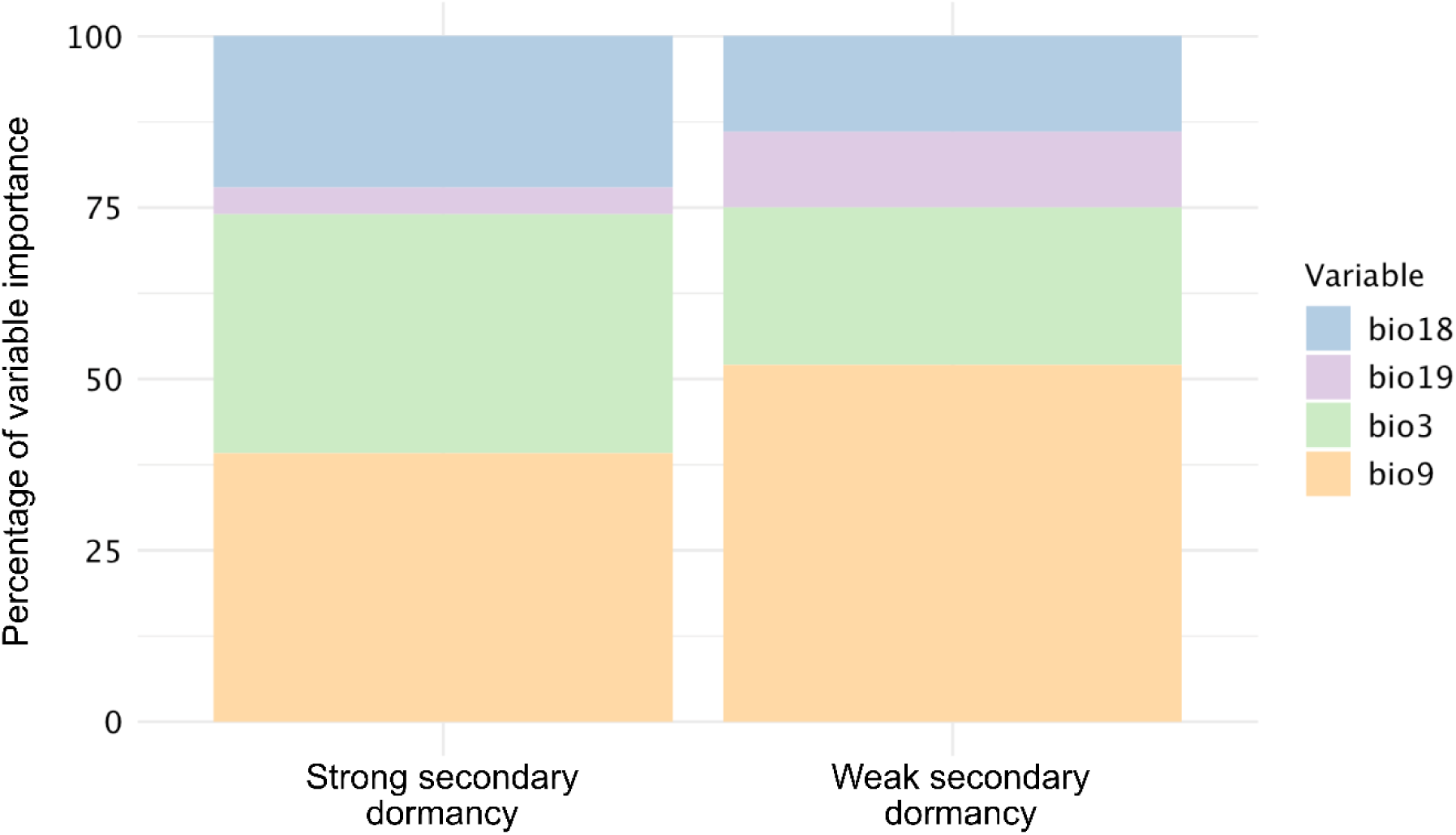
Importance of predictor variables in the species distribution model for strong and weak heat-induced secondary dormancy ecotypes. The relative importance of four bioclimatic variables in the species distribution model (SDM) used to predict the habitat suitability for strong and weak secondary dormancy ecotypes of Arabidopsis thaliana. The variables included in the model are isothermality (BIO3), mean temperature of the driest quarter (BIO9), mean precipitation of the warmest quarter (BIO18), and mean precipitation of the coldest quarter (BIO19). Trial 1 data were used for this analysis.

**Figure S6.**
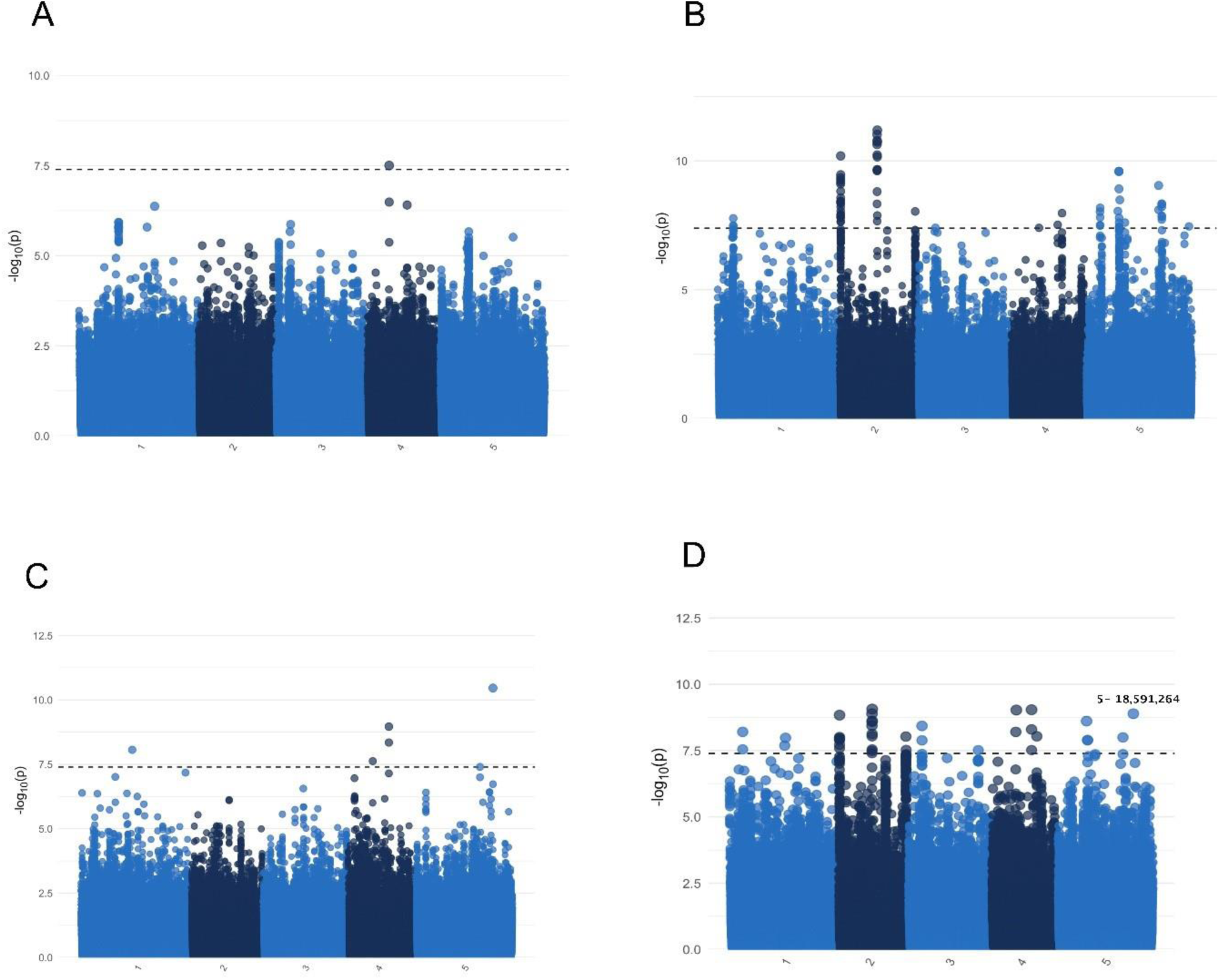
Manhattan plot displaying the association of 1.2M SNP markers with primary dormancy across trials. This plot presents the genome-wide association results for primary dormancy using 1.2 million SNP markers and three independent replicates of European Arabidopsis thaliana genotypes. (A) Results for Trial 1 of 295 lines, (B) Trial 2 of 361 lines, and (C) Trial 3 of 344 lines. Population structure was controlled for in the analysis using a kinship matrix between individuals. (D) The combined results from all three trials were calculated using Fisher’s combined P-value method. A significant peak on chromosome 5 in panel (D) is annotated, which is located just 14 bp from the region of the DELAY OF GERMINATION 1 (DOG1) gene, a well-known regulator of seed dormancy. The dashed horizontal line indicates the significance threshold after a 5% Bonferroni correction was applied.

**Figure S7.**
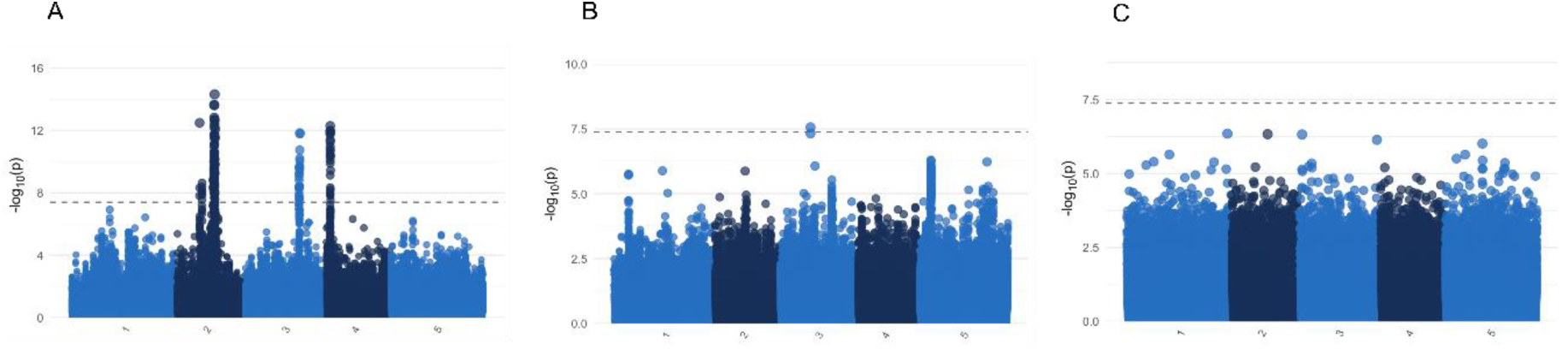
Manhattan plot displaying the association of 1.2M SNP markers with heat-induced secondary dormancy across trials. The genome-wide association results for heat-induced secondary dormancy using 1.2 million SNP markers across three independent replicates of European Arabidopsis thaliana genotypes. The analysis includes 295 lines in Trial 1 (A), 361 lines in Trial 2 (B), and 344 lines in Trial 3 (C). Primary dormancy was included as a covariate in the analysis, and population structure was accounted for using a kinship matrix between individuals. The dashed horizontal line represents the significance threshold after a 5% Bonferroni correction was applied. Significant loci are highlighted and indicate genomic regions associated with heat-induced secondary dormancy. So-called sdorm treatment represents germination rates following a 3-day stratification at 4°C to release dormancy, followed by a 4-day treatment at 37°C. The pdorm treatment tested germination rates without any pre-treatment, and the control treatment involved seeds stratified for 3 days at 4°C to release dormancy before testing germination. All germination tests were conducted under long-day conditions at 20°C, and germination rates were recorded after 7 days.

## Tables

**Table S1.** Information of 361 studied accessions. Table records Genotype ID (GenotypeID) according to ENA database and Genotype Name (GenotypeName) accordingly. Geographical origin (origin) and precise latitude and longitude of the genotype are recorded, together with extra information such as sequencer, collector, and CSS accession ID (AccessionID). Attached as a separate file.

**Table S2.** Spearman correlation coefficients for heat-induced secondary dormancy across three trials. The Spearman correlation values quantifying the consistency of heat-induced secondary dormancy measurements across three independent trials. The secondary dormancy treatment represents germination rates following a 3-day stratification at 4°C to release dormancy, followed by a 4-day treatment at 37°C. The germination test was conducted under long-day conditions at 20°C, and germination rates were recorded after 7 days. All germination tests were conducted under long-day conditions at 20°C, and germination rates were recorded after 7 days. Trial 1 included 295 genotypes and was conducted in May 2022; Trial 2 comprised the full set of 361 genotypes and was conducted in December 2022; Trial 3 included 344 genotypes and was conducted in December 2023.

|  | rho | p value |
| --- | --- | --- |
| Trial 1 vs Trial 2 | 0.494 | < 2.2e-16 |
| Trial 2 vs Trial 3 | 0.409 | 3.661e-12 |
| Trial 1 vs Trail 3 | 0.350 | 4.393e-09 |

**Table S3.**
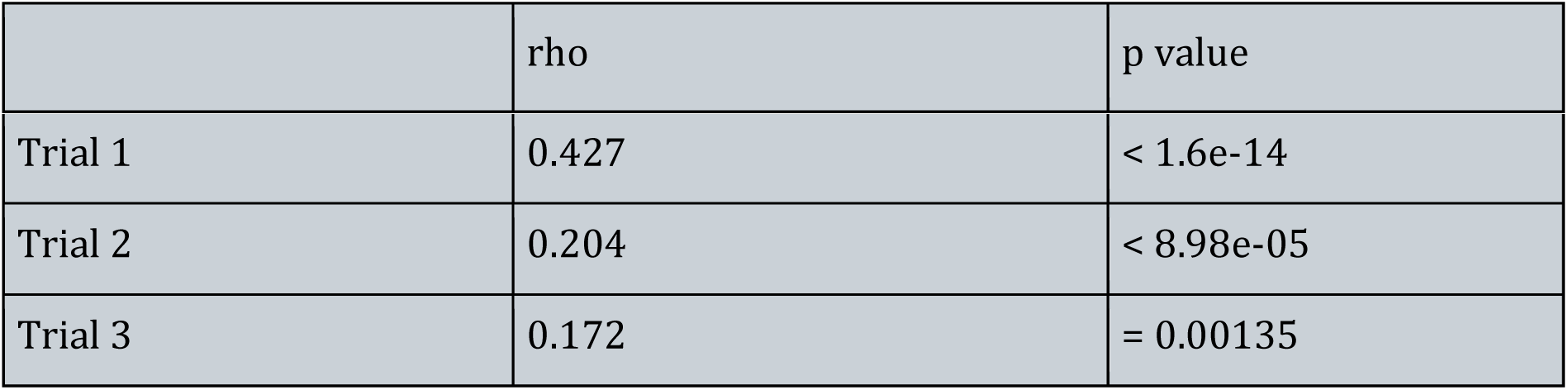
Spearman correlation of residual primary dormancy and heat-induced secondary dormancy of three trials. Secondary dormancy treatment represents germination rates following a 3-day stratification at 4°C to release dormancy, followed by a 4-day treatment at 37°C. The primary dormancy treatment tested germination rates without any pre-treatment. All germination tests were conducted under long-day conditions at 20°C, and germination rates were recorded after 7 days. Trial 1 is a set of 295 samples, Trial 2 has the complete set of 361 samples, and Trial 3 is a set of 344 samples. Trial 1 was performed in May 2022, Trial 2 in December 2022, and Trial 3 in December 2023.

|  | rho | p value |
| --- | --- | --- |
| Trial 1 | 0.427 | < 1.6e-14 |
| Trial 2 | 0.204 | < 8.98e-05 |
| Trial 3 | 0.172 | = 0.00135 |

**Table S4.** Descriptive statistics of germination rates under heat-induced secondary dormancy treatment across low-(<50°) and high-latitude (≥50°) regions in three trials. We used 50oN as a threshold to separate northern and southern populations based on our assumptions of the biogeographical transitions in Central and Northern Europe. Secondary dormancy treatment represents germination rates following a 3-day stratification at 4°C to release dormancy, followed by a 4-day treatment at 37°C. All germination tests were conducted under long-day conditions at 20°C, and germination rates were recorded after 7 days. Trial 1 is a set of 295 samples, Trial 2 has the complete set of 361 samples, and Trial 3 is a set of 344 samples. Trial 1 was performed in May 2022, Trial 2 in December 2022, and Trial 3 in December 2023.

|  | Low latitude | High latitude |
| --- | --- | --- |
| Trial 1 | mean = 0.0511, var = 0.0278 | mean = 0.1808, var = 0.0826 |
| Trial 2 | mean = 0.1999, var = 0.1012 | mean = 0.4193, var = 0.152 |
| Trial 3 | mean = 0.2546, var = 0.0611 | mean = 0.3871, var = 0.0502 |

**Table S5.** Regression results of heat-induced secondary dormancy with four bioclimatic variables as predictors of genetic variation in germination after three treatments, across three trials. The results of regression models that assess the influence of four bioclimatic variables on secondary dormancy and its genetic variation. The models employed a binomial likelihood with a logit link function. The bioclimatic variables included BIO3 (isothermality), BIO9 (mean temperature of the driest quarter), BIO18 (mean precipitation of the warmest quarter), and BIO19 (mean precipitation of the coldest quarter). The analysis was performed across three treatments: primary dormancy (pdorm), secondary dormancy (sdorm), and control. Secondary dormancy was induced by a 4-day 37°C treatment following a 3-day stratification at 4°C to release primary dormancy. Primary dormancy was tested without any pre-treatment, whereas control seeds were stratified at 4°C for 3 days before germination testing. The sdorm value used in this model represents secondary dormancy corrected for primary dormancy, calculated as the residual of the regression of secondary dormancy on primary dormancy. All germination tests were conducted under long-day conditions at 20°C, and germination rates were recorded after 7 days. Trial 1 is a set of 295 samples, Trial 2 has the complete set of 361 samples, and Trial 3 is a set of 344 samples. Trial 1 was performed in May 2022, Trial 2 in December 2022, and Trial 3 in December 2023. (A), (B), and (C) represent models with the control group as the baseline for Trial 1, Trial 2, and Trial 3, respectively; (D), (E), and (F) represent models in whichprimary dormancy is used as the baseline, for Trial 1, Trial 2, and Trial 3, respectively. Attached as a separate file.

**Table S6.** Genome-wide association results of primary dormancy across three trials and shared genome-wide association peaks across three experimental trials. The primary dormancy treatment tested germination rates without any pre-treatment. All germination tests were conducted under long-day conditions at 20°C, and germination rates were recorded after 7 days. (A) Trial 1 is a set of 295 samples, (B) Trial 2 is the complete set of 361 samples, and (C) Trial 3 is a set of 344 samples. Trial 1 was performed in May 2022, Trial 2 in December 2022, and Trial 3 in December 2023. (D) Shared genome-wide association peaks of primary dormancy across all experimental trials, computed by Fisher’s combined probability test. Attached as a separate file.

**Table S7.** Genome-wide association results of heat-induced secondary dormancy across three trials and shared genome-wide association peaks across three experimental trials. The secondary dormancy represents germination rates following a 3-day stratification at 4°C to release dormancy, followed by a 4-day treatment at 37°C. All germination test s were conducted under long-day conditions at 20°C, and germination rates were recorded after 7 days. (A) Trial 1 is a set of 295 samples, (B) Trial 2 is the complete set of 361 samples, and (C) Trial 3 is a set of 344 samples. Trial 1 was performed in May 2022, Trial 2 in December 2022, and Trial 3 in December 2023. (D) Shared genome-wide association peaks of heat-induced secondary dormancy across all experimental trials, computed by Fisher’s combined probability test. Attached as a separate file.

**Table S8.** Definitions of bioclimatic (BIO) variables used in the analysis and interpretation of their values. Descriptions of the nineteen bioclimatic (BIO) variables included in the generalized linear mixed model (GLMM). The ecological meaning of each BIO variable is provided, along with guidance on interpreting their values—particularly clarifying conditions represented by low and high values.

| Bioclimatic variables | Definition | Intepretation |
| --- | --- | --- |
| BIO1<br>Annual Mean Temperature | Mean of monthly<br>temperature averages | High: warmer overall<br>climate; Low: cooler year-round |
| BIO2<br>Mean Diurnal Range | Mean of monthly (max<br>temp - min temp) | High: large day-night<br>temperature shifts |
| BIO3<br>Isothermality | $= (\text{BIO2} / \text{BIO7}) \times 100$ | High: more uniform temp<br>across year (diurnal ≈<br>seasonal); Low: high<br>seasonal contrast relative<br>to daily variation |
| BIO4<br>Temperature Seasonality | Standard deviation of<br>temperature $\times 100$ | High: large seasonal<br>variability; Low: stable<br>seasonal temps |
| BIO5<br>Max Temperature of<br>Warmest Month | Highest monthly max<br>temperature | High: hot extremes in<br>summer |
| BIO6<br>Min Temperature of<br>Coldest Month | Lowest monthly min<br>temperature | Low: cold winters |
| BIO7<br>Temperature Annual<br>Range | $= \text{BIO5} - \text{BIO6}$ | High: strong summer-<br>winter contrast; Low: mild<br>year-round |
| BIO8<br>Mean Temperature of<br>Wettest Quarter | Average temp during 3-month wettest period | Ecologically varies;<br>important for timing of<br>growth or dormancy |
| BIO9<br>Mean Temperature of<br>Driest Quarter | Average temp during 3-month driest period | High: warm/dry periods;<br>Low: cold/dry stress |
| BIO10<br>Mean Temp of Warmest Quarter | Average temp during warmest 3 months | High: hot growing seasons |
| BIO11<br>Mean Temp of Coldest Quarter | Average temp during coldest 3 months | Low: cold stress potential |
| BIO12<br>Annual Precipitation | Total yearly precipitation | High: wet climates; Low: arid regions |
| BIO13<br>Precipitation of Wettest Month | Total precipitation in the wettest month | High: strong rainy season |
| BIO14<br>Precipitation of Driest Month | Total precipitation in driest month | Low: drought severity |
| BIO15<br>Precipitation Seasonality | Coefficient of variation of monthly precipitation | High: unpredictable rain patterns |
| BIO16<br>Precipitation of Wettest Quarter | Precipitation total in wettest 3-month period | High: monsoon-heavy systems |
| BIO17<br>Precipitation of Driest Quarter | Precipitation total in driest 3-month period | Low: drought-prone periods |
| BIO18<br>Precipitation of Warmest Quarter | Precipitation in the warmest 3 months | Low: hot/dry stress; High: wet growing season |
| BIO19<br>Precipitation of Coldest Quarter | Precipitation in coldest 3 months | Low: potential for overwintering stress |

